# Optimizing Phylogenetic Eigenvector Regression: Union Eigenvectors, Robust Estimation, and Flexible Application to Comparative Analyses

**DOI:** 10.1101/2024.04.14.589420

**Authors:** Zheng-Lin Chen, Deng-Ke Niu

**Author notes:** E-mail addresses (Z.-L. Chen), (D.-K. Niu). Correspondence author: Deng-Ke Niu, College of Life Sciences, Beijing Normal University, Beijing, 100875, China.

## Abstract

Phylogenetic eigenvector regression (PVR) is widely used in ecology and evolution by representing phylogenetic structure through separable eigenvectors. Despite this flexibility, its implementation faces three key challenges: (1) the selection of eigenvectors, (2) the reduced robustness of ordinary least-squares (OLS) regression under shift-like evolutionary heterogeneity, and (3) the applicability of conventional model complexity rules such as the "samples-per-variable (SPV) ≥ 10" guideline. Here, we propose an optimized PVR framework that addresses these limitations. First, we show that trait-specific selections of eigenvectors often diverge, sometimes producing inconsistent results, and that using their union offers stronger control of phylogenetic non-independence. Second, we evaluate robust regression estimators within PVR, demonstrating that PVR-MM – and in most cases PVR-L2, the standard OLS estimator – maintains high accuracy under non-stationary evolutionary shifts where other non-robust methods fail. Third, through simulation, we reassess the SPV ≥ 10 rule, showing that PVR tolerates eigenvector counts well beyond this threshold, offering greater flexibility while requiring attention to potential overfitting. Extensive simulations across diverse trees and evolutionary scenarios confirm that the optimized framework improves accuracy and robustness. By addressing key aspects of eigenvector selection, regression, and model complexity, our findings strengthen the reliability and applicability of PVR.

## Introduction

Comparative studies of traits across species must account for phylogenetic non-independence (Cornwallis & Griffin, 2024; Dewar et al., 2025; Felsenstein, 1985; Garamszegi, 2014). Among the available approaches, phylogenetic eigenvector regression (PVR) has become widely used in ecology and evolutionary biology because it expresses phylogenetic information as separable eigenvectors (EVs) obtained by decomposing the phylogenetic distance or covariance structure encoded in the tree (Diniz-Filho et al., 2012a; Diniz-Filho et al., 2014; Diniz-Filho et al., 2011; Diniz-Filho et al., 1998; Diniz-Filho & Tôrres, 2002). This representation provides flexibility because it captures phylogenetic structure directly from the tree without requiring the specification of an explicit evolutionary model, while also accommodating extensions to other forms of autocorrelation such as spatial or temporal structure (Kühn et al., 2009; Maynard et al., 2022; Peres-Neto et al., 2012; Safi & Pettorelli, 2010; Wang et al., 2019) and with machine learning methods (Debastiani et al., 2021; Penone et al., 2014). Such versatility has helped promote the broad adoption of PVR in ecological applications and highlights the importance of refining its implementation (Divíšek et al., 2018; Gao et al., 2023; Lanuza et al., 2023; LeRoy et al., 2020).

Despite its utility, two issues limit PVR performance. First, EVs selected from each trait separately may yield inconsistent results, raising concerns when analyzing relationships between traits. Second, conventional PVR relies on ordinary least squares regression, which can be sensitive to non-stationary evolutionary shifts or outliers. These challenges motivate the development of optimized strategies that improve reliability while retaining the flexibility of PVR.

Here, we propose an optimized PVR framework that combines (i) a union-based EV set (*EV_U*) to capture shared phylogenetic structure, (ii) robust regression estimators to reduce sensitivity to non-stationarity (Adams et al., 2024; Huber, 1964; Slater & Pennell, 2014; Yu & Yao, 2017), and (iii) a reassessment of the conventional "samples-per-variable (SPV, i.e., events per variable) ≥ 10" rule (Babyak, 2004), showing that PVR remains reliable with far fewer samples per predictor and emphasizing the need to check EV counts before model fitting. Using extensive simulations under a range of evolutionary scenarios and tree structures, we assess the accuracy of these strategies relative to established comparative methods and highlight the conditions under which optimized PVR offers particular advantages.

## Materials and Methods

### Simulation of phylogenetic trees

We evaluated PVR on a set of fully bifurcating trees chosen to vary in branch-length scaling, taxon sampling, and topological balance, generated with *stree* (**ape** 5.8) and, where necessary, combined with *paste.tree* (**phytools** 2.3-0) (Paradis & Schliep, 2019; Revell, 2024):

1. **Baseline**: 128-tip balanced tree, Grafen (1989) scaling ρ = 1. (ρ is the exponent in Grafen’s branch-length transformation; ρ = 1 yields equal branch-length scaling across hierarchical levels.)
2. **Compressed deep branches**: 128-tip balanced tree, ρ = 0.1 (short ancestral, long terminal branches).
3. **Elongated deep branches**: 128-tip balanced tree, ρ = 2 (long ancestral, short terminal branches).
4. **Extreme imbalance**: 128-tip ladder (comb-shaped) tree, ρ = 1.
5. **Small-sample setting**: 16-tip tree generated under the same four branch-length conditions (balanced with ρ = 1, 0.1, 2, and ladder).

### Simulated trait generation

We generated paired traits *X*_1_ and *X*_2_ on each of the eight phylogenetic trees under six evolutionary settings adapted from Revell (2010), Adams et al. (2024), and Chen et al. (2025). In all scenarios, the generative model for *X*_2_ is *X*_2_ = *βX*_1_ + *ε*, where *β* denotes the regression slope specifying the intended linear dependence of *X*_2_ on *X*_1_, and *ε* contributes additional noise (Table 1).

**Table 1.** Six evolutionary scenarios for simulating paired traits *X*_1_ and *X*_2_ (generated as *X*_2_ = *βX*_1_ + *ε*).

| Scenario | $X_1$ | $\varepsilon$ | Shift? | $\beta$ | Explanation/purpose |
| --- | --- | --- | --- | --- | --- |
| Gradual-evolution scenarios (stationary) |  |  |  |  |  |
| BM–BM | BM | BM | No | 1 | $X_1$ and $\varepsilon$ both follow BM, so $X_2$ is a linear combination of two BM components. As a result, realized associations between $X_1$ and $X_2$ can vary substantially across replicates, particularly as stochastic variance increases, despite a fixed structural relationship in the data-generating process. |
| BM–Norm | BM | Normal | No | 1 | $X_1$ follows BM, while $\varepsilon$ is drawn from an i.i.d. normal (non-phylogenetic) distribution. The non-phylogenetic $\varepsilon$ reduces the phylogenetic signal in $X_2$, causing realized associations to vary across replicates and, under high noise variance, potentially weaken the correlation relative to that implied by the generative $\beta$. |
| Norm–BM | Normal | BM | No | 1 | $X_1$ is i.i.d. normal; $\varepsilon$ adds phylogenetic structure to $X_2$. Realized correlations between $X_1$ and $X_2$ vary across replicates and may be substantially attenuated by noise relative to that implied by the generative $\beta$. |
| Norm–Norm | Normal | Normal | No | 1 | Neither trait carries phylogenetic signal. $X_1$ and $X_2$ vary independently, and realized associations may be weak or highly variable depending on noise variance. |
| Shift scenarios (non-stationary, single-branch abrupt shifts) |  |  |  |  |  |
| $X_1$-shift | BM + shift | Normal | $X_1$ only | 1 | Extends the BM–Norm scenario by adding a single-branch shift to $X_1$, while fixing $\varepsilon \sim \mathcal{N}(0, 0.01)$. Because $\varepsilon$ is very small, the background evolutionary association between $X_1$ and $X_2$ remains tightly coupled across most of the phylogeny, whereas the shifted branch introduces a localized deviation from this otherwise strong relationship. |
| Dual-shift | BM + shift | BM + shift | Both | 0 | $X_1$ and $\varepsilon$ both undergo a shift on the same branch, and no causal association is intended between $X_1$ and $X_2$. Both $X_1$ and $\varepsilon$ are simulated independently under BM, with the same root value (1) and variance ($\sigma^2 = 1$), ensuring that shifts occur on the same scale. Such synchronized shifts may induce spurious overall correlation signals in methods that lack robustness to localized shifts. |

The generative model for $X_2$ consists of a linear combination of $X_1$ and random noise $\varepsilon$ , with $\beta$ controlling the intended association. In the two shift scenarios, an additional shift parameter $s$ displaces trait means on a single deep branch. A series of values ( $10^{-4}$ , 0.01, 1, 4, 16, 64, 256, 1024) was used to model both the shift magnitude and the gradient of $\varepsilon$ 's variance, influencing either the variance of $\varepsilon$ in stationary models or the shift size in non-stationary scenarios, and thereby modulating the strength of population-level correlations. Brownian-motion (BM) traits were simulated using *rTraitCont* (**ape** 5.8) with rate = 1, and normal values were generated using *rnorm* (**stats** 4.1.3). Each magnitude was replicated 1,000 times, yielding 8,000 datasets per scenario and 48,000 datasets per tree. For implementation details of the shift simulations, see Chen et al. (2025).

These six settings fall into two broad categories: (i) four gradual-evolution scenarios, in which traits evolve without abrupt changes, and (ii) two shift scenarios, in which an abrupt change is introduced on a single deep branch. The four gradual-evolution scenarios (Table 1) involve traits with noise that either evolved under Brownian motion (BM) or with independent and identically distributed (i.i.d.) normal (non-phylogenetic) noise (Norm). In all scenarios, *X*_1_ is generated first, and *X*_2_ is then constructed as a linear function of *X*_1_ plus a noise term *ε*. The four gradual-evolution scenarios differ in whether *X*_1_ and *ε* evolve under Brownian motion (BM) or independent normal noise (Norm), yielding the combinations BM–BM, BM–Norm, Norm–BM, and Norm–Norm. For example, in the BM–Norm scenario, *X*_1_ follows Brownian motion, whereas *X*_2_ inherits the phylogenetic structure of *X*_1_ through the *βX*_1_ term, with additional independent and identically distributed (i.i.d.) normal noise *ε*. The variance of *ε* is determined by a numerical gradient (10⁻⁴, 0.01, 1, 4, 16, 64, 256, 1024), modifying noise intensity and phylogenetic dilution, here referring to the progressive weakening of phylogenetic signal in trait correlations as non-phylogenetic noise increases. In contrast, in the two shift scenarios (*X*_1_-shift and Dual-shift, Table 1), the same numerical gradient specifies the shift magnitude *s* applied to one of the two sister branches descending directly from the root, producing non-stationary, abrupt evolutionary changes in *X*_1_ alone or in both *X*_1_ and *ε*, where non-stationary refers to localized deviations from homogeneous evolutionary dynamics along the phylogeny. Together, these six scenarios span a wide range of phylogenetic signals, noise intensities, and stationary versus non-stationary dynamics, providing a robust and diverse framework for benchmarking PVR variants and comparison methods. For clarity, the full combination of evolutionary scenarios, variance or shift magnitudes, and replicate counts is summarized in Table 1.

### Defining process-level ground truth

A commonly used approach in comparative-method simulations is to define true and false positives directly by the generating effect size *β*, with *β* = 0 representing an absence of true association and *β* ≠ 0 representing a true association (Adams & Collyer, 2022). This strategy is intuitive and effective for simulation designs in which the structural signal specified by *β* is expressed consistently across datasets, and where stochastic variation and finite sample effects do not substantially alter the resulting data-level association patterns.

However, our simulation framework differs in a key respect: because trait evolution is stochastic on a fixed phylogeny, each replicate expresses its own realized statistical association, determined jointly by the coupling parameter *β* and the stochastic variation encoded by the noise component *ε*. As a result, the association observed in any given dataset may diverge from that implied by the generative value of *β*.

Most notably, when *β* =1, the noise term *ε* can be sufficiently large – particularly under higher-variance settings such as 256 or 1024 – to mask or overwhelm the *β*-induced linear trend. In such cases, the realized population-level association may be weak, often leading to nonsignificant statistical results in finite samples despite a nonzero generative *β*. By the same logic, when *β* = 0, any apparent association reflects stochastic evolutionary variation and may be observed in individual realizations despite the absence of structural coupling.

Consequently, a method’s "true" performance cannot be assessed by whether it recovers statistical relationship implied by the coupling parameter *β*; instead, evaluating performance requires a dataset-specific benchmark that reflects the realized association in each replicate.

To accomplish this, we used branchwise standardized changes (*ΔX*/*L*, where *L* denotes branch length) as a replicate-specific, phylogenetically independent benchmark. For every simulation, trait values at all nodes were recorded and standardized changes were computed on each branch (*ΔX*_1_/*L* and *ΔX*_2_/*L*, for each trait respectively). Because these branchwise changes are independent across the tree, they provide a tree-wide ground-truth sample for the statistical association realized in that dataset. This branchwise differencing approach is conceptually related to Felsenstein’s (1985) phylogenetically independent contrasts (PIC), which also leverage the independence of evolutionary changes along branches. But unlike PIC, our procedure does not require ancestral-state reconstruction or any Brownian-motion assumptions, as all node states are known in our simulations.

We quantified the realized association using Spearman’s rank correlation across the branchwise changes, which provides a robust, rank-based summary of the association realized in each dataset. We then evaluated each comparative method against this benchmark: a method was considered correct only if it recovered both (i) the direction of association and (ii) the significance decision (*p* < 0.05) for that replicate. Across 1,000 replicates for each experimental condition, method accuracy was defined as the proportion of correct inferences.

### Phylogenetic eigenvectors selection methods

EVs were obtained by principal-coordinate analysis of the double-centered cophenetic distance matrix, yielding *n* − 1 axes. For each trait (*X*_1_ or *X*_2_), we selected a subset of EVs with three published criteria (Diniz-Filho et al., 2012a; Diniz-Filho & Nabout, 2009):

1. **ESRRV**: keep EVs whose Pearson or Spearman correlation with the focal trait is significant, choosing the test according to the trait’s normality.
2. **mAIC**: forward stepwise regression adding EVs that lower Akaike Information Criterion (AIC), implemented with *stepAIC()* (**MASS** 7.3-55) (Venables & Ripley, 2002).
3. **mMorI**: forward selection until the residual Moran’s I is minimized. Because phylogenetic autocorrelation is almost always positive, we modified this rule to stop as soon as residual Moran’s I becomes negative; if it never does, selection continues to the usual non-significant threshold.

Custom R wrappers *ESRRV()* and *MoransI()* implement rules 1 and 3, while all other calculations use the original **PVR** package (Diniz-Filho et al., 2012a; Diniz-Filho et al., 1998; Diniz-Filho et al., 2012b).

### Correcting phylogenetic autocorrelation in trait regressions

After EV selection we compared four ways to purge phylogenetic signal:

1. ***EV_X*_1_ model**: *X*_2_ ∼ *X*_1_ + *EV_ X*_1_ — covariates are the EVs chosen for *X*_1_.
2. ***EV_X*_2_ model**: *X*_2_ ∼ *X*_1_ + *EV_ X*_2_ — uses the EVs chosen for *X*_2_. Writing it in reverse form, *X*_1_ ∼ *X*_2_ + *EV_ X*_2_, controls the same EVs, so the correlation (and its *p*-value) is identical, although the regression slope becomes the reciprocal of the original.
3. ***EV_U* model**: *X*_2_ ∼ *X*_1_ + *EV_U* — *EV_U* is the union of EVs selected from both traits.
4. **Residual correlation**: Regress each trait on its own EVs, then take the Pearson correlation of the residuals. To curb type-I error from over-fitting, *p*-values were based on *df* = *n* – *k* – 1, where *n* is the number of species and *k* the number of EVs in *EV*_*U*. While this conservative adjustment may reduce power, it controls for inflated false positives due to shared structure among EVs (Diniz-Filho & Tôrres, 2002).

### Robust regressions in PVR

To test the sensitivity of PVR to outliers and non-normal errors we compared it with five different regression estimators:

1. **L2**: as termed in robust regression (Huber, 1964), corresponds to the standard ordinary least-squares (OLS) estimator that minimizes squared residuals under homoscedastic normal errors. It was implemented using *lm()* (package **stats**, R 4.1.3).
2. **L1**: least-absolute-deviation (*lad()*, **L1pack** 0.41-245) (Phillips, 2002).
3. **M**: Huber-type M-estimator (*rlm()*, **MASS** 7.3-55) (Venables & Ripley, 2002).
4. **S**: high-breakdown S-estimator (*lmRob()*, **robust** 0.7-4) (Gervini & Yohai, 2002).
5. **MM**: S start followed by an M step (*lmRob()*, **robust**).

Among these estimators, L2 serves as the baseline; L1, M, S, and MM offer progressively stronger resistance to leverage points while retaining efficiency. For detailed properties and phylogenetic applications of these estimators, see Adams et al. (2024).

## Results

Unless otherwise noted, in the subsequent analyses, PVR regressions control phylogeny by including as covariates the EVs selected for each trait separately (*EV_ X*_1_, *EV_ X*_2_) or their union (*EV_U*).

### Eigenvector selection and budgeting

We applied three EV-selection rules (ESRRV, mAIC, mMorI) separately to *X*_1_ and *X*_2_; the resulting EV sets, together with their union (*EV*_*U*), were carried forward as candidate covariates for examination. As detailed later through mathematical derivations and PVR accuracy assessments in the section “*Why use the union of eigenvectors?*”, *EV*_*U* plays an important role in controlling phylogenetic autocorrelation, so here we give special attention to ∣*EV*_*U*∣ (the number of eigenvectors in EV_U) when comparing outcomes across rules, without disregarding the other EV sets.

Across all four tree scenarios with 128 species, the size of the *EV*_*U* set differed markedly among selection rules (Fig. 1). mAIC consistently returned the largest sets, with medians ranging from 123 to 125 EVs across topologies. By comparison, ESRRV produced *EV_U* medians ranging from 6 to 31, and mMorI from 1 to 37; under the balanced 128-species tree with ρ =1, both methods returned a median of 9.

**Fig. 1.**
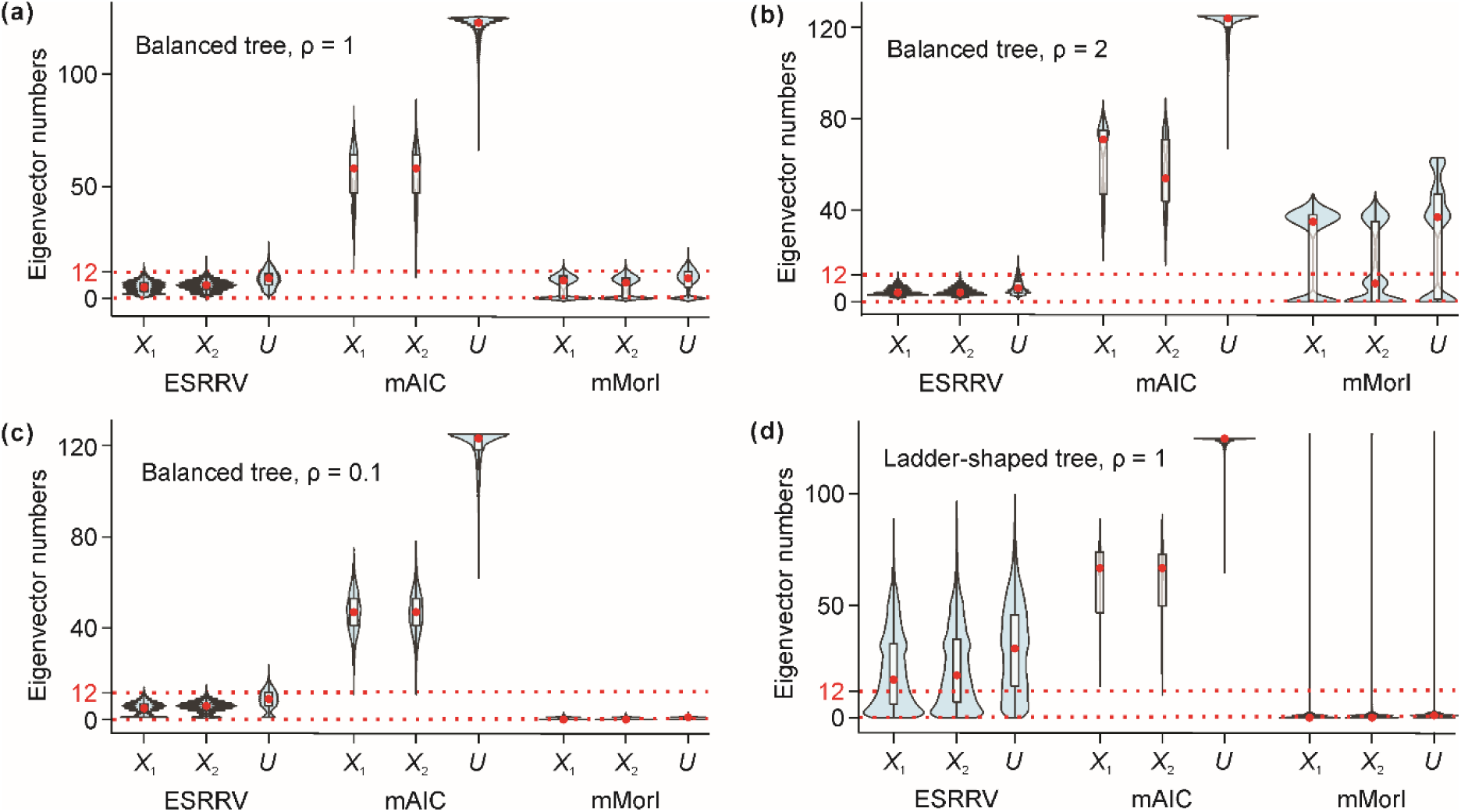
Number of eigenvectors (EVs) retained by three selection criteria (ESRRV, mAIC and mMorI) for a 128-species phylogenetic tree. EVs were picked separately for *X*_1_ and *X*_2_; *EV*_*U* is their union. Violin outlines depict the full distribution of selected EV counts; white boxes show the inter-quartile range, black whiskers the 5th–95th percentiles, and red dots mark the median of each distribution. The red dashed line marks the upper bound implied by the conventional samples-per-variable ≥ 10 guideline, calculated here assuming one non-EV predictor (128 spp./10 = 12.8, meaning 11.8 EVs plus 1 predictor). Selection criteria are described in Materials and Methods.

The four scenarios with 16 species (Fig. S1) showed the same relative ordering across selection rules and tree types, even though the absolute *EV_U* counts scaled downward proportionally with the reduced tree size. This confirms that EV counts are influenced primarily by tree topology and the selection rule, rather than sample size.

In our SPV calculations, the sample size is the number of species at the tips of the tree, and the variable count includes both the retained EVs and the single predictor trait (*X*_1_), giving |EV| + 1 variables in total. The widely recommended "SPV ≥ 10" rule (Austin et al., 2017; Babyak, 2004; Peduzzi et al., 1996) originally proposed to prevent over-parameterization and unstable inference in conventional regression – states that SPV should be at least 10. So for our 128 species trees, this would imply an upper bound of 12.8 variables (i.e., ∼12 EVs + 1 predictor). However, because EVs are orthogonal components derived from the phylogenetic distance matrix rather than biological or environmental predictors, this rule does not carry the same meaning in PVR as it does in ecological regressions.

To test this idea, we analyzed the subset of simulated datasets corresponding to balanced trees with ρ = 1 and *n* = 128 species, using six evolutionary scenarios. For each scenario, we examined five |*EV*_*U*| settings, corresponding to SPV ratios of approximately 1.5, 2, 3, 6, and the EV count selected by mMorI. In this setting, mMorI typically returned fewer EVs than the conventional SPV ≥ 10 threshold yet was already sufficient to control phylogenetic autocorrelation and avoid false positives due to residual structure. We then augmented the mMorI-selected *EV_U* core set (median |EV_U| = 9) by adding additional EVs in descending eigenvalue order to reach the 6, 3, 2, and 1.5 SPV ratio targets (|EV_U| = 20, 42, 63, and 84, respectively). When independence was effectively addressed by the EV selected using mMorI, differences in PVR accuracy across these *EV_U*-count gradients are attributable mainly to potential overfitting from excessive predictors.

In the six simulation scenarios, the impact of EV count on PVR accuracy varied. Here ’PVR accuracy’ denotes the accuracy of the PVR-controlled trait–trait regression *X*_2_ ∼ *X*_1_ + *EV*_*U*, as defined in the Materials and Methods, where the benchmark for determining correct inference is described in detail.

Across the four gradual-evolution scenarios (Figs. 2a–d), accuracy showed distinct patterns as EV count increased. In the three gradual-evolution cases (Figs. 2a, 2b, and 2d), accuracy began to decline when |*EV_U*| = 63 and declined further at |*EV_U*| = 84. In the Norm–BM scenario (Fig. 2c), accuracy exhibited only modest variation across EV counts, and the mMorI-selected set (median = 9 EVs) was not among the best-performing choices.

**Fig. 2.**
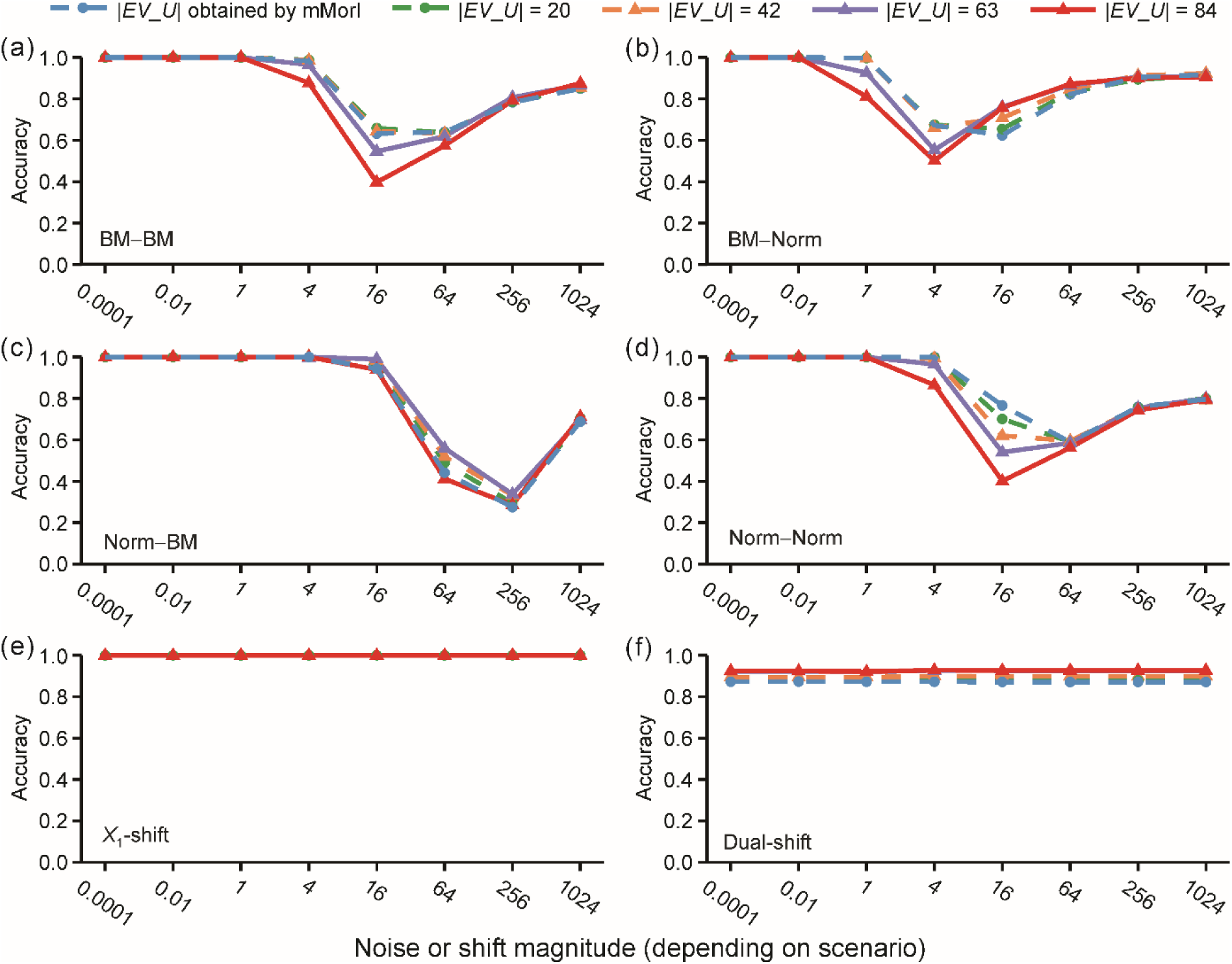
Accuracy of PVR under six simulation scenarios with varying numbers of eigenvectors (*EV_U*). Panels (a–d) show the four gradual-evolution (stationary) scenarios, whereas panels (e–f) show the two shift scenarios involving a single-branch abrupt evolutionary change. All simulated on a balanced phylogenetic tree with 128 species and uniformly scaled branch lengths (ρ = 1). Lines represent PVR accuracy for different *EV_U* sizes. The mMorI-selected *EV_U* (median |*EV_U*| = 9) serves as the core set, and larger *EV_U* sizes (|*EV_U*| = 20, 42, 63, and 84) were obtained by augmenting this core set with additional eigenvectors in descending eigenvalue order. Accuracy is defined with respect to each dataset’s realized association, based on replicate-specific branchwise standardized changes (*ΔX*/*L*). For details on EV selection, scenario and accuracy definitions, see Materials and Methods; the data used to plot this figure appear in Table S1.

We next turn to the two non-stationary scenarios (Figs. 2e–f), which introduce an abrupt evolutionary shift on a deep branch, either in *X*_1_ alone or in both *X*_1_ and *ε*. Under the single-shift condition (Fig. 2e), accuracy was nearly identical across all EV counts, indicating no sensitivity to EV number. In the dual-trait shift scenario (Fig. 2f), accuracy differed only slightly across EV counts. Unlike Figs. 2a, 2b and 2d – where the *EV_U* obtained directly by mMorI was among the strongest choices and modest EV increases performed similarly – the larger EV sets (63 and 84) showed a marginal but consistent accuracy advantage over the *EV_U* set obtained directly by mMorI.

Taken together, these results in Fig. 2 show that modest increases in EV count – such as expanding from the mMorI-selected set (median = 9 EVs) to intermediate sizes around 40 EVs, corresponding to a SPV ratio of roughly 3 – do not dramatically reduce accuracy under gradual-evolution scenarios, and under shift scenarios the PVR results are more insensitive to EV number. These patterns reflect the behavior of EVs themselves rather than the performance of any particular EV-selection strategy evaluated in Fig. 1 and Fig. S1, and also illustrate why the conventional SPV-based heuristic does not directly translate to PVR, given the orthogonality of phylogenetic eigenvectors.

These findings also clarify the implications for EV-selection rules. Across all tree topologies (Fig. 1 and Fig. S1), mAIC consistently returned very large EV sets, often far exceeding the range in which accuracy stabilizes in Fig. 2, and is therefore unsuitable for PVR. ESRRV generally produced moderate EV counts and can be applied across topologies, but mMorI frequently returned very few EVs – including zero in some unbalanced trees – which limits its ability to control phylogenetic autocorrelation. However, from a methodological perspective, mMorI has the conceptual advantage of explicitly minimizing residual Moran’s I – the most direct criterion for removing phylogenetic dependence. Accordingly, in balanced trees with ρ = 1, where ESRRV and mMorI return similar EV counts, the comparative results presented in subsequent sections indicate that mMorI may be preferable because it directly optimizes phylogenetic residual structure.

### Trait-specific selection of EVs and its downstream effects on PVR estimation

Although retaining somewhat larger EV sets does not markedly reduce PVR accuracy overall, most empirical applications of PVR select eigenvectors only for the predictor variable, implicitly assuming that the EV sets selected for different traits do not differ enough to influence inference (e.g., Álamo et al., 2024; Carotenuto et al., 2015; Gao et al., 2023; Liu et al., 2024; Seger et al., 2025; Vilela et al., 2014). However, whether the EV sets selected for different traits coincide or diverge – and, if they diverge, whether such differences materially affect PVR estimation – remains unclear. This motivates a systematic evaluation of trait-specific selection of EVs and its consequences.

To explore this, we conducted simulations using balanced phylogenetic trees with 128 species and uniformly scaled branches (ρ = 1). We analyzed paired predictors under various evolutionary scenarios to quantify how divergence in EV sets impacts regression outcomes.

Across 8,000 simulations per scenario (Table 2), the EV sets selected for two traits (*X*_1_ and *X*_2_) disagreed in 76.4% of cases on average. The highest rate of disagreement occurred under the dual-trait shift scenario, with notable inconsistencies observed across all scenarios. These results indicate that EV selection is highly sensitive to which trait is treated as the dependent variable.

**Table 2.** Frequency with which the EV sets chosen for *X*_1_ and *X*_2_ do not match.

| Simulation scenarios | ESRRV | mMorI |
| --- | --- | --- |
| BM–BM | 83.3% | 84.1% |
| BM–Norm | 80.4% | 81.4% |
| Norm–BM | 86.3% | 75.8% |
| Norm–Norm | 86.0% | 4.5% |
| $X_1$-shift | 91.0% | 49.2% |
| Dual-shift | 94.5% | 100% |

Given the high disagreement rates between the EV sets selected for *X*_1_ and *X*_2_ (Table 2), we next examined whether these differences translate into divergent PVR outcomes. To do so, we fit parallel PVR models using either *EV*_*X*_1_ or *EV_X*_2_ as covariates and compared the resulting inferences on the *X*_1_–*X*_2_ relationship. Table 3 reports the proportion of replicates in which the two regressions yielded different conclusions (i.e., differing in direction or significance). These analyses quantify the practical consequences of EV-set discrepancies: when *X*_1_ and *X*_2_ select different eigenvectors, the downstream PVR conclusions may also diverge. The magnitude of this effect varies across evolutionary scenarios, with the largest discrepancies occurring under the BM–BM and dual-shift conditions when EVs were selected by ESRRV.

**Table 3.** Proportion of simulations in which PVR models using the EV set from *X*_1_ versus that from *X*_2_ yield discordant results.

| Simulation scenarios | ESRRV | mMorI |
| --- | --- | --- |
| BM–BM | 19.8% | 11.4% |
| BM–Norm | 9.18% | 18.2% |
| Norm–BM | 11.5% | 10.8% |
| Norm–Norm | 6.44% | 0.13% |
| $X_1$-shift | 14.1% | 6.02% |
| Dual-shift | 30.7% | 14.9% |

To further quantify the extent of divergence, we computed the absolute difference in partial correlation coefficients for the discordant cases:

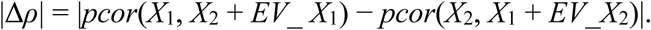

Over 73% of discordant comparisons showed |Δ*ρ*| > 0.1, and 15% exceeded 0.5, demonstrating substantial variation in effect size and even sign reversals in biological interpretation (Fig. 3). The *X*_1_-only shift scenario was especially extreme, with median |Δ*ρ*| values of 1.04 (ESRRV) and 0.97 (mMorI). Even among the 73,318 cases where models agreed in significance and slope sign, 31% showed |Δ*ρ*| > 0.1, and 4% exceeded 0.5 (Fig. S2). Thus, consistent significance does not necessarily imply consistent effect estimation.

**Fig. 3.**
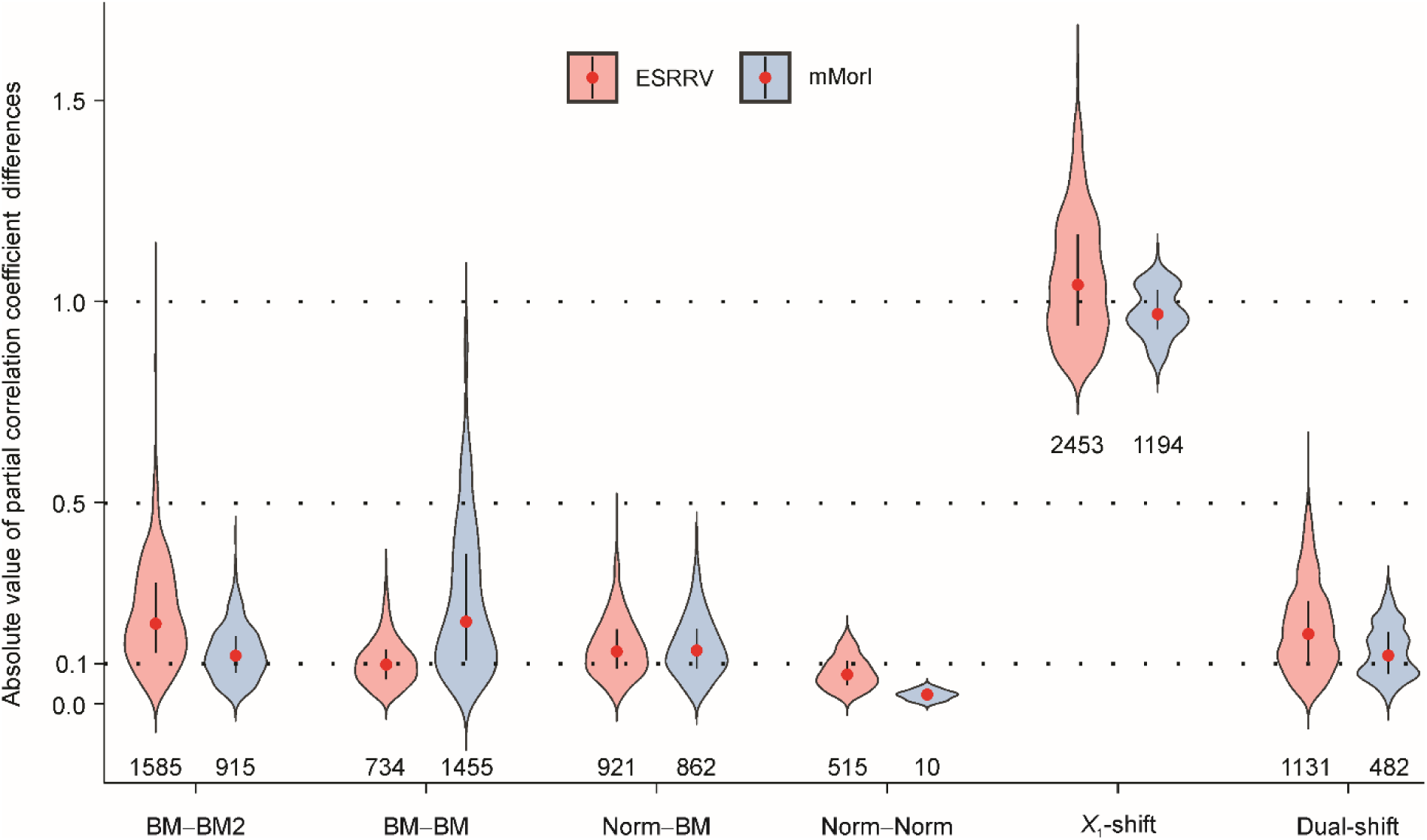
Distributions of absolute differences in partial correlation coefficients between comparing PVR models using *EV*_*X*_1_ versus *EV*_*X*_2_ as covariates under six evolutionary scenarios. The violin plot illustrates the absolute differences in partial correlation coefficients |pcor(*X*_1_, *X*_2_ + *EV*_*X*_1_) − pcor(*X*_2_, *X*_1_ + *EV*_*X*_2_)| for cases with conflicting PVR results; numbers beneath each violin give the sample size (*n*). Partial correlations were computed with ppcor::pcor() (Pearson). Red dots indicate the median values, black vertical bars span the inter-quartile range, and black whiskers extend to the 5th–95th percentiles. Paired t-tests revealed significant differences in partial-correlation estimates between *EV*_*X*_1_ and *EV*_*X*_2_ in every scenario (all *p* < 0.001) except when mMorI was applied to the Norm−Norm scenario data, where the test was non-significant (*n* = 10, *p* = 0.373). See Materials and Methods for details of the EV-selection procedures (ESRRV, mMorI) and the BM, Norm and Shift simulation variants; the raw values plotted here are provided in Table S4.

In summary, trait-specific selections of EVs frequently diverge (Table 2), and in a measurable subset of cases these divergences lead to different PVR conclusions, manifested as differences in significance or moderate-to-large coefficient shifts. The magnitude of these effects depends on the evolutionary context, with certain scenarios (e.g., dual-shift or trait-specific shifts) showing particularly pronounced instability. Moreover, preliminary results (Fig. 1b) indicate that under alternative branch-length scaling (ρ = 2), the EV sets selected for *X*_1_ versus *X*_2_ – when under the mMorI rule – differ far more than in the baseline ρ = 1 setting (Fig. 1a), suggesting an even greater potential for distorted inference.

### Why use the union of eigenvectors?

The EV sets selected for *X*_1_ and *X*_2_ frequently differ (Table 2), and these differences lead to distinct PVR outcomes in a subset of replicates (Table 3). This inconsistency prompts a more fundamental theoretical question: can such per-trait EV sets adequately control for phylogeny when two traits share only part of their phylogenetic signal?

When either trait is filtered through only its own eigenvector set, phylogenetic structure unique to the partner trait remains unmodeled, potentially biasing the estimated association. Our derivation (Appendix A) shows that using the union of both EV sets (*EV*_*U*) more fully captures the phylogenetic component (P) in Cheverud’s (1985) decomposition of phenotypic variation (T = P + S), resulting in more stable and unbiased regression estimates. This aligns with Westoby et al.’s (2023) critique that phylogenetic generalised least squares (PGLS) corrects for phylogeny in the response but not in the predictor.

Motivated by the same concern, we additionally considered a two-step alternative: each trait is first regressed on its own EVs to remove trait-specific phylogenetic signal, and then an OLS regression is applied to the resulting residuals to quantify their association. This residual-based approach offers a different path to adjusting for phylogeny in both variables and provides a comparative baseline for evaluating the *EV_U* method.

Building on the earlier results (Fig. 1a), where both ESRRV and mMorI produced admissible EV counts under a balanced tree with 128 species and ρ = 1, we next used simulations to assess whether the choice of selection method materially influences PVR outcomes. Additionally, we tested whether regressions based on *EV_U* outperform those relying on EVs selected separately for each trait or on residual approaches.

Figure 4 compares eight PVR configurations: two EV-selection methods (ESRRV, mMorI), each combined with four regression strategies (*EV*_*X*_1_, *EV*_*X*_2_, *EV*_*U*, and residual). Because accuracy is averaged over 1,000 replicates – and EV-set differences affect outcomes only in a subset of cases (Table 3) – the four regression strategies appear broadly similar. When *EV*_*U* equals the larger of *EV*_*X*_1_ or *EV*_*X*_2_, the corresponding curves overlap, further reducing visual differences among methods. Despite this overall similarity, systematic differences still emerge. Across scenarios, *EV_U*-based regressions (red lines) – regardless of whether paired with ESRRV (dashed) or mMorI (solid) – generally perform better than the other regression strategies (blue, green, and black lines). Among these, mMorI + *EV_U* (red solid line) consistently ranked first or jointly first across all six evolutionary scenarios and shift magnitudes, making it the most reliable configuration overall.

**Fig. 4.**
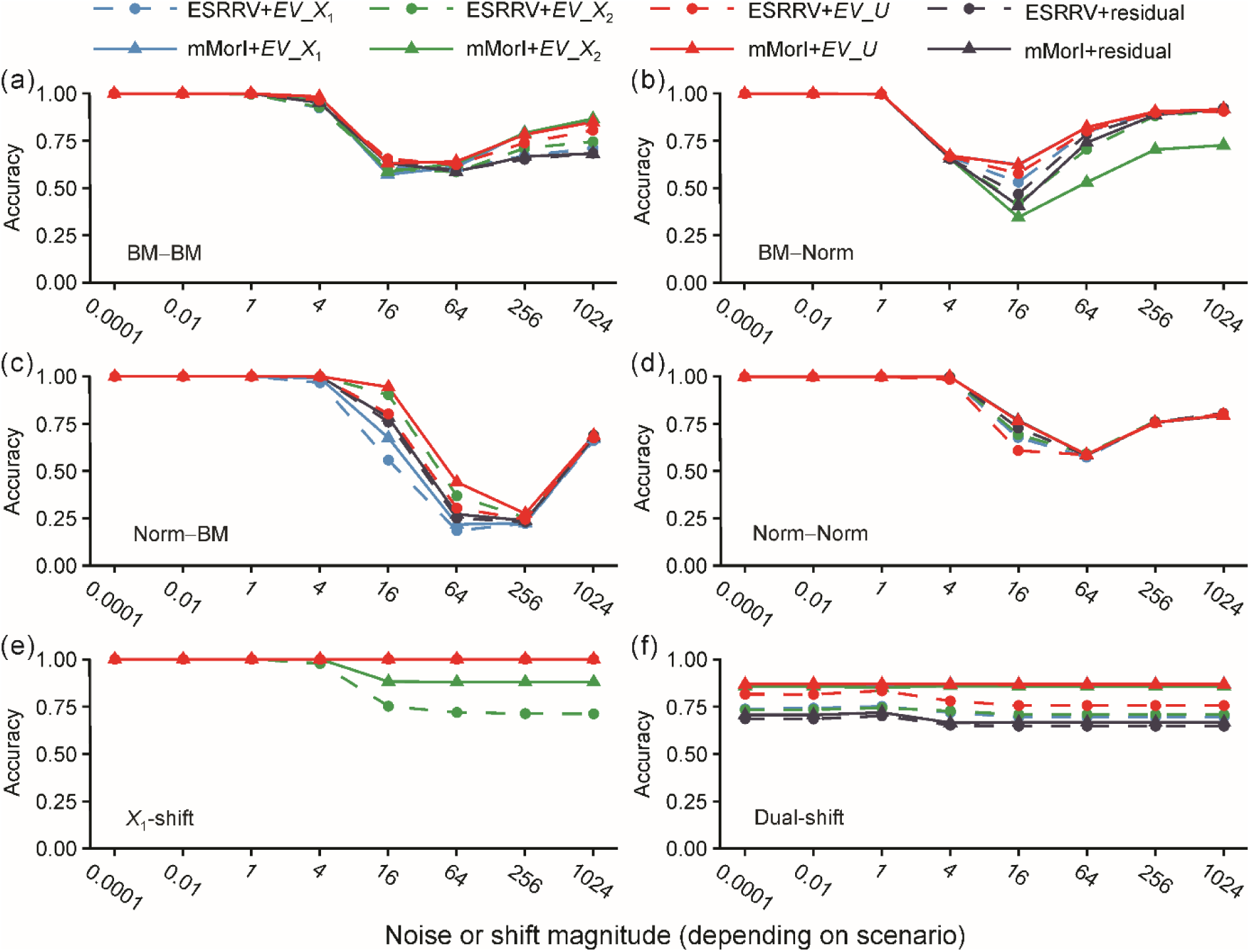
Performance across different phylogenetic eigenvector (EV) selection methods and regression strategies. This figure compares two EV selection methods (ESRRV and mMorI), each combined with four regression approaches: (1) using *EV*_*X*_1_ as covariates; (2) using *EV*_*X*_2_; (3) using their union, *EV_U*; and (4) regressing the residuals of *X*_1_ and *X*_2_, each pre-adjusted by their respective EVs. Simulations were conducted on balanced trees with 128 species (ρ = 1) across six scenarios: (a–d) the four gradual-evolution (stationary) scenarios – BM–BM, BM–Norm, Norm–BM, and Norm–Norm – and (e–f) the two single-branch shift scenarios – *X*_1_-only shift and dual-trait shift. In some cases, when multiple method combinations produced identical results, their lines overlap and may appear as a single curve in the plot. Accuracy values are evaluated against each dataset’s realized association, estimated from replicate-specific branchwise standardized changes. For details on EV selection, scenario and accuracy definitions, see Materials and Methods. The source data for this figure are provided in Table S5.

While residual-based regressions (black lines) also account for phylogenetic signals in both traits, their performance was generally inferior to that of *EV_U*-based strategies (Fig. 4). This discrepancy likely arises because removing phylogenetic signal independently for each trait can also eliminate shared structure that underlies true biological associations. In contrast, *EV_U* integrates information from both traits within a single model, thereby preserving shared structure while controlling for phylogenetic effects, and consequently achieved higher accuracy across scenarios.

A notable special case occurs in Fig. 4e, where the two traits are highly correlated but *X*_1_ undergoes a large, localized shift on a deep branch. Under this condition, all configurations – except those relying on *EV*_*X*_2_ – yield nearly identical results. The overlapping lines illustrate this convergence among *EV_U*, *EV*_*X*_1_, and residual approaches, likely reflecting the fact that they each capture a critical subset of eigenvectors that effectively model the key phylogenetic signal introduced by the shift in *X*_1_.

Across scenarios where both ESRRV and mMorI produced admissible EV counts, we retained mMorI as the default selection method, owing to its consistent superiority. When considered together with results from Fig. 1 and Fig. 2, our findings suggest that mMorI is preferable for balanced trees with ρ = 1 or ρ = 2, while ESRRV should be used for ρ = 0.1 balanced trees and ρ = 1 ladder-shaped trees, where mMorI often fails to return sufficient EVs.

### EV_U-based PVR as an intrinsically robust framework

We evaluated five regression estimators – L2 (ordinary least squares), L1 (least absolute deviations), S (high-breakdown S-estimator), M (Huber-type M-estimator), and MM (S-start followed by an M-step) – within the *EV_U*-based PVR framework. Throughout this section, L2-based PVR is used as the baseline for comparison. To assess robustness across evolutionary contexts, we focused on the combinations of tree topology and EV-selection method identified in the preceding analyses. Specifically, this analysis included 128-species trees with four phylogenetic topologies: balanced trees with ρ = 1 and ρ = 2 analyzed under mMorI, and balanced trees with ρ = 0.1 and ladder-shaped trees with ρ = 1 analyzed under ESRRV (Figs. S3–S6). In addition, 16-species balanced trees with ρ = 1 were analyzed under mMorI (Fig. S7). Across these settings, L2, M, and MM yielded nearly identical accuracies in most scenarios, with their curves largely overlapping in the plots. Only in ladder-shaped trees under strong shift conditions did MM show a clear robustness advantage, while L2 and M remained indistinguishable. Compared with these three estimators, the performance of L1 varied depending on the scenario; in contrast, S generally underperformed across multiple shift settings and was thus excluded from subsequent analyses. Estimator M was also omitted from later figures, as its outputs were nearly indistinguishable from L2 and were almost entirely overlaping in all scenarios, making its inclusion redundant.

### Comparative evaluation of optimized PVR, PGLS, and PIC

To compare optimized PVR with established phylogenetic regression methods, we evaluated multiple PVR configurations alongside PGLS and PIC-based approaches. Here, L2 serves as the standard least-squares baseline, whereas L1 and MM represent increasingly robust alternatives. We used *EV_U* constructed from mMorI for 128-species balanced trees (ρ = 1, 2) and 16-species balanced trees (ρ = 1), and from ESRRV for 128-species balanced trees (ρ = 0.1) and ladder-shaped trees (ρ = 1). These approaches were compared against three established methods, PGLS, PIC-L2 (i.e., PIC-OLS), and PIC-MM (a robust variant recommended by Adams et al. (2024). For PGLS, the best-fitting evolutionary model was selected from five candidates – BM, Pagel’s λ, Ornstein–Uhlenbeck (OU) with fixed α, OU with random α, and the Early-Burst (EB) model – using AIC and is hereafter referred to as PGLS-optimal.

Across all phylogenetic structures examined, the four gradual-evolution scenarios (BM–BM, BM–Norm, Norm–BM, and Norm–Norm) produced broadly similar accuracy patterns among most methods (Figs. 5a–d; Figs. S8–S11a–d). In these settings, the various PVR variants generally performed slightly below – or occasionally comparable to – PGLS-optimal and PIC-L2, and often showed accuracy similar to PIC-MM. Under the BM–Norm condition, however, all methods except PGLS-optimal performed similarly, with most showing somewhat higher accuracy than PGLS-optimal. These similarities across figures reflect that, under gradual or near-gradual evolutionary dynamics, the compared phylogenetic regression approaches capture the major components of phylogenetic structure to a comparable degree.

**Fig. 5.**
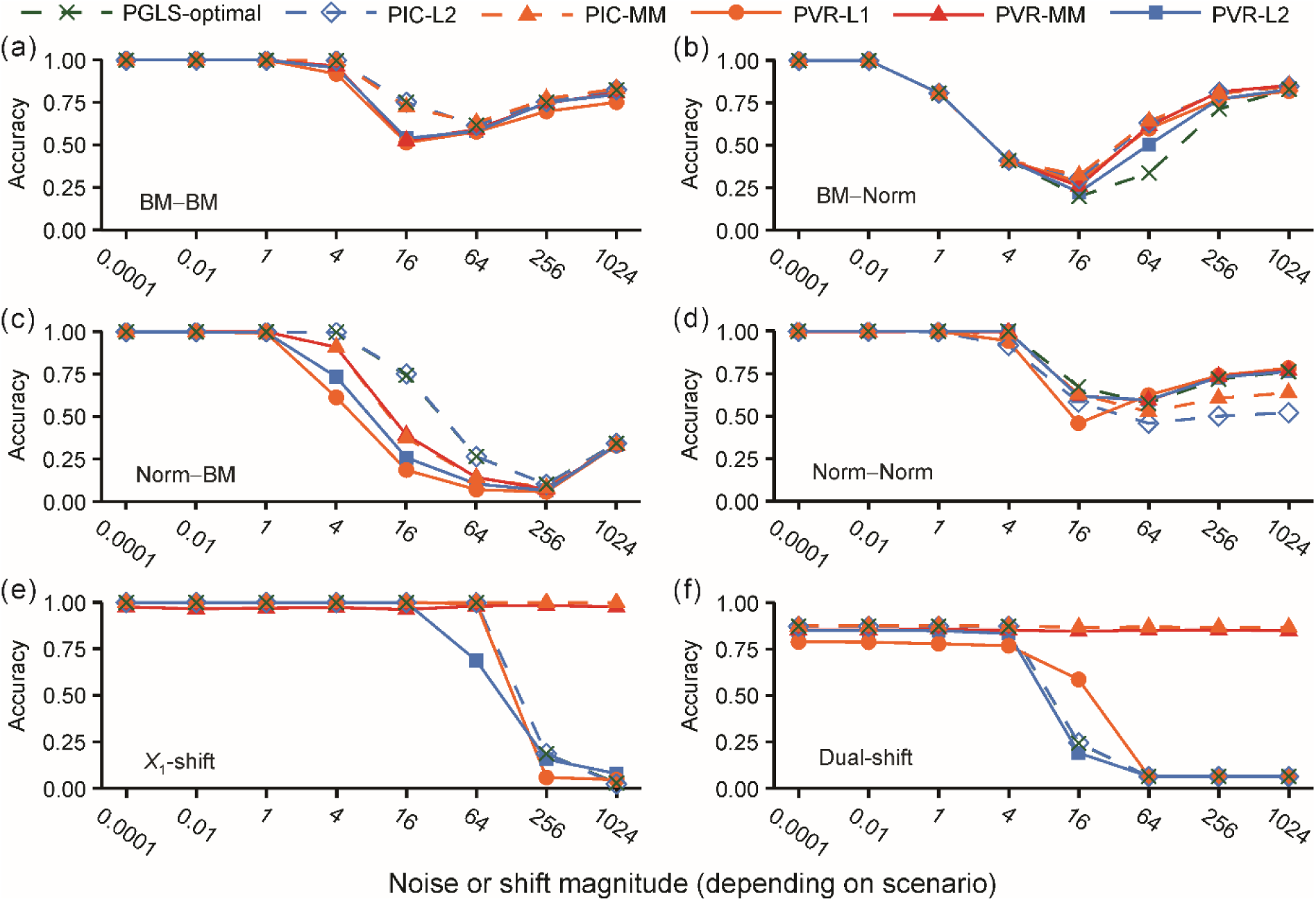
Performance of PVR, PGLS, and PIC on 128-species ladder-shaped phylogenetic trees (ρ = 1). Comparisons were conducted under six evolutionary scenarios: (a–d) the four gradual-evolution (stationary) scenarios – BM–BM, BM–Norm, Norm–BM, and Norm–Norm – and (e–f) the two non-stationary single-branch shift scenarios – *X*_1_-only shift and dual-trait shift. All eigenvectors (EVs) were selected using the ESRRV criterion. Accuracy is quantified using a replicate-level benchmark derived from branchwise standardized changes. For details on ESRRV, scenario definitions, and accuracy measures, see Materials and Methods. The data underlying this figure are provided in Table S6.

In contrast, the shift scenarios revealed clearer performance differences (Figs. 5e–f; Figs. S8–S11e–f). Across all tree shapes – whether 128-species or 16-species, and across balanced trees with varying values of ρ – PGLS-optimal and PIC-L2 consistently showed poor robustness when shift magnitudes increased. By comparison, PIC-MM and the PVR variants remained markedly more robust to strong evolutionary shifts, with even the standard PVR-L2 regression showing substantial robustness under most tree shapes (Figs. S8–S11e–f). On ladder-shaped trees (Figs. 5e-f), however, the robustness of PVR-L1 and PVR-L2 deteriorated, yielding performance closer to that of PGLS-optimal and PIC-L2. In this setting, only the MM-based approaches – PIC-MM and PVR-MM – sustained high accuracy and effectively accommodated large shifts.

## Discussion

PVR, first proposed to model trait variation and trait–trait correlations using phylogenetic eigenvectors (Diniz-Filho et al., 1998), is widely used across ecological studies. Its strength lies in capturing separable, orthogonal eigenvectors, without requiring an explicit evolutionary model. This flexibility allows easy incorporation into machine learning methods such as random forests and neural networks (Debastiani et al., 2021; Penone et al., 2014), supporting tasks like trait imputation and classification beyond regression. Moreover, because the eigenvectors are orthogonal, they can be jointly modeled with other sources of autocorrelation (e.g., spatial or temporal), a flexibility particularly valuable in ecological settings where multiple forms of structure coexist (Kühn et al., 2009; Maynard et al., 2022; Safi and Pettorelli, 2010; Wang et al., 2019).

### Methodological improvements to PVR

In this study, we revisit and optimize the PVR framework. We show that eigenvectors derived from different traits often diverge, and using their union (*EV_U*) more effectively controls phylogenetic non-independence. While our focus is on trait pairs, the logic extends to multi-trait settings, where incorporating all relevant eigenvectors selected for each trait helps retain phylogenetic signal. However, care must be taken to avoid overfitting by limiting the number of retained eigenvectors.

The "SPV ≥ 10" rule is widely recommended in conventional regression to guard against overfitting and unstable inference (Austin et al., 2017; Babyak, 2004; Peduzzi et al., 1996). Because eigenvectors are orthogonal rather than correlated predictors, uncritically importing such heuristics into phylogenetic contexts risks misrepresenting model reliability. Our analyses indicate that PVR remains reliable with much lower SPV, under gradual evolution, with accuracy still acceptable when SPV falls to ∼3, whereas under shift conditions PVR performance shows virtually no such lower bound. This tolerance affords greater flexibility, crucial for small datasets: with ≤19 species, SPV ≥ 10 precludes eigenvectors entirely, whereas a relaxed threshold of SPV ≥ 3 allows inclusion of up to four eigenvectors. Our analyses further suggest a practical guideline: regardless of the selection method, researchers should inspect the number of retained eigenvectors before incorporating them into regression models. For example, mAIC typically returns excessively large sets that risk overfitting, whereas mMorI may frequently yield very few or even zero eigenvectors.

### PVR under non-stationary and shift-like evolutionary dynamics

Unlike likelihood-based methods such as PGLS, whose performance is inherently tied to a single evolutionary model (Freckleton et al., 2011), PVR extracts phylogenetic structure through eigenvector decomposition and thus remains flexible when traits deviate from standard processes (e.g., BM, OU, EB) or when model uncertainty is high. Because phylogenetic eigenvectors are derived solely from the topology and branch-length structure of the tree, they capture variation expressed at both broad and fine phylogenetic scales. This property makes them well suited for describing complex empirical patterns – such as clade-specific differences in trait lability, heterogeneous evolutionary rates, or multiple adaptive regimes – that cannot be adequately summarized by a single global parameter (Diniz-Filho et al., 2015; Diniz-Filho et al., 2014; Diniz-Filho et al., 2019; Diniz-Filho et al., 2012b; Diniz-Filho et al., 2010; Gouveia et al., 2013). Within this perspective, the abrupt evolutionary shifts examined in our simulations represent an extreme form of non-stationarity: substantial localized departures from Brownian covariance that arise when evolutionary rates or selective regimes change along particular branches of the tree. With appropriately selected EV sets (e.g., union-based selection guided by mMorI or ESRRV), EVs can capture this structure, yielding near-i.i.d. residuals. Because M-estimator reweighting is primarily triggered by influential outliers (Huber, 1964), it is rarely activated under such conditions; consequently, default PVR-L2 (OLS) typically matches PVR-M in robustness, outperforming PGLS or typical PIC. Under extreme heterogeneity (e.g., strongly ladderized trees with multiple sharp shifts), PVR-MM affords added stability. Rather than overturning prior assessments characterizing PVR as inferior to likelihood-based approaches (Freckleton et al., 2011; Laurin, 2010; Pennell & Harmon, 2013), our results build upon earlier demonstrations that PVR’s performance depends critically on how eigenvectors are selected, and show that systematic selection strategies can substantially improve stability and accuracy under non-stationary evolutionary conditions (Diniz-Filho et al., 2015; Diniz-Filho et al., 2012a; Diniz-Filho et al., 2010).

### PGLS under signal imbalance and unmodeled phylogenetic structure

Our analyses reveal that the performance of PGLS is strongly shaped by how phylogenetic signal is distributed between the two traits. In Fig. 5 and Figs. S8–S11, PGLS-optimal performed worst in the BM–Norm condition, where the dependent variable (*X*_2_) carried markedly weaker phylogenetic signal than the predictor (*X*_1_). In contrast, PGLS achieved the best or near-best performance in the other stationary conditions, in which *X*_2_ carried substantially stronger or at least comparable signal. This pattern is consistent with the fact that PGLS inherits from OLS the assumption that predictors are fixed and measured without error (Casson & Farmer, 2014; Jarantow et al., 2023; McArdle, 1988; Osborne & Waters, 2002), creating a mismatch when the predictor carries the stronger phylogenetic signal. Preliminary analyses by Chen et al. (2023, preprint) similarly show that such signal imbalance can distort regression estimates, underscoring the importance of accounting for the distribution of phylogenetic signal when selecting the response variable.

Adding phylogenetic eigenvectors to PGLS – a framework recently explored by Schraiber et al. (2024) could, in principle, also mitigate problems arising when the predictor carries substantially stronger phylogenetic signal than the response, as in the BM–Norm scenario. Although Schraiber et al. did not study this mechanism, their empirical results highlight a related phenomenon. In fungal gene-expression data, including eigenvectors in PGLS markedly reduced spurious correlations, revealing unmodeled phylogenetic structure that is unlikely to reflect differences in gradual signal strength between paired genes. Instead, their results point to abrupt, localized deviations from Brownian expectations – a shift-like form of non-stationarity closely paralleling the abrupt dual-shifts examined in our simulations. These findings illustrate a broader function of eigenvectors: they can capture residual phylogenetic structure not encoded in the assumed covariance matrix, whether arising from gradual imbalance or from localized evolutionary shifts. Within this broader context, determining which eigenvectors should be added to PGLS remains an open question. A natural possibility – yet to be formally evaluated – is that including only those eigenvectors present in the predictor but absent in the response may provide a better complement to PGLS than using *EV_U*, or perhaps even using the full set of predictor-selected eigenvectors, including those that overlap with the response. Because PGLS already accounts for the phylogenetic structure of the response through its covariance model, re-introducing eigenvectors that primarily reflect that same structure could risk unnecessary complexity or overfitting. Exploring whether predictor-specific eigenvectors yield more stable inference therefore represents a promising direction for future work.

### Extending asymmetry considerations to spatial autocorrelation models

Although our focus is on phylogenetic comparative methods, similar issues arise in spatial analyses, where spatial autocorrelation must also be accounted for (Cottenie, 2005; Dale & Fortin, 2009; Diniz-Filho & Bini, 2005; He et al., 2003; Keitt et al., 2002; Legendre, 1993; Peres-Neto & Legendre, 2010). Preliminary tests show that the asymmetry introduced by variable designation – discussed for PVR – also affects spatial frameworks such as spatial eigenvector mapping, CAR, SAR, and trend-surface GAMs, with performance differing when dependent and independent variables are swapped (Table S7; see Spatial_methods in Supplementary material for implementation details). By analogy, the union-based strategy we adopted for PVR may also apply to spatial analyses, potentially helping to preserve shared spatial signal while controlling autocorrelation. In addition, our findings regarding effective sample sizes – particularly the observation that reliable performance can be achieved with substantially lower SPV than conventionally assumed – may also be informative for spatial models, where limited sample sizes and large sets of spatial eigenvectors pose similar challenges.

### Concluding remarks: Practical guidelines for selecting eigenvectors in PVR

Based on our theoretical derivations and extensive simulations across evolutionary scenarios and tree structures, we recommend the following practical guidelines for selecting eigenvectors in PVR:

1. **Use the union of eigenvectors selected separately for each trait as the default.** This strategy consistently outperforms per-trait selections by controlling phylogenetic structure in both variables and preventing asymmetric corrections.
2. **Prefer mMorI when it does not return too few eigenvectors; otherwise use ESRRV.** mMorI typically provides a compact set of eigenvectors by minimizing residual Moran’s I, ensuring strong phylogenetic correction. However, when mMorI selects very few – or even zero – eigenvectors, ESRRV offers a more reliable alternative by providing a more adequate number of components.
3. **Avoid mAIC for regression-based PVR**, because the resulting EV sets are typically excessively large, risking overfitting without improving phylogenetic control.
4. **Inspect EV counts before model fitting.** For datasets with *n* ≤ 19, SPV ≥ 10 would prohibit the use of eigenvectors entirely. However, our simulations show that PVR remains reliable with SPV ≈ 3, while shift-like non-stationarity allows an even more relaxed SPV threshold.
5. **Prefer PVR under non-stationary or shift-like evolutionary conditions.** Under these scenarios, PVR maintains strong performance even without robust estimation. However, under more extreme heterogeneity – such as strongly ladderized trees with sharp shifts – using a robust MM-estimator (PVR-MM) can provide additional stability.

## Supplementary material

Supplementary material is submitted with this manuscript as online Supporting Information.

## Data availability

The scripts used to generate the results in the main text—including phylogenetic tree construction, trait data simulation, regression model fitting, and statistical analysis—are provided in Main_code.zip. The code used to evaluate spatial statistical methods is provided in Spatial_methods.zip. In addition, we developed an R package, ROBPVRU, which implements phylogenetic eigenvector regression with union operations on eigenvectors combined with robust estimators. All code and data used to generate Fig. 1 and the supplementary figures have been uploaded in a review-only format to preserve author anonymity during peer review and will be made freely accessible via the first author’s GitHub page upon acceptance.

