## Supplementary text and figures for "Optimizing Phylogenetic Eigenvector Regression: Union Eigenvectors, Robust Estimation, and Flexible Application to Comparative Analyses"

Text S1.

**Theoretical Justification for Using the Union of Eigenvectors Selected for Each Trait**

We derive a multiple regression framework based on the decomposition model T = P + S (Cheverud et al., 1985), using two traits denoted as *X*₁ and *X*₂ (e.g., body weight and metabolic rate) without assuming a fixed causal direction, to illustrate how phylogenetic signals from both traits can influence regression outcomes.

**1. Decomposition of Two Traits**
Let *X*₁ and *X*₂ represent two continuous traits. For both traits:
  T = P + S
  *X*₁ = P₁ + S₁, *X*₂ = P₂ + S₂

**2. Partitioning the Phylogenetic Component**
Assume both traits share part of the phylogenetic signal (EV₁), and each also has unique structure:
  P₁ = *α*₁·*EV*₁ + α₂·*EV*₂
  P₂ = *α*₃·*EV*₁ + α₄·*EV*₃

**3. Modeling the Trait Relationship**
We model S₂ as a function of S₁:
  S₂ = *β*·S₁ + *ε*
Substituting in the trait model:
  *X*₂ – P₂ = *β*·(*X*₁ – *P*₁) + *ε*
Rewriting:
  *X*₂ = *β*·*X*₁ + (P₂ – *β*·P₁) + *ε*

**4. Expanding the P Terms and Simplifying**
Substitute P₁ and P₂:
  *X*₂ = *β·X*₁ + (*α*₃·*EV*₁ + *α*₄·*EV*₃ – *β·α*₁·*EV*₁ – *β·α*₂·*EV*₂) + *ε*
Define:
  *β*₁ = (*α*₃ – *β·α*₁), *β*₂ = –*β·α*₂, *β*₃ = *α*₄
Then:
  *X*₂ = *β·X*₁ + *β*₁·*EV*₁ + *β*₂·*EV*₂ + *β*₃·*EV*₃ + *ε*

This equation shows that the relationship between traits is confounded by phylogenetic structure from both traits.

**Conclusion**
This derivation demonstrates that using only the eigenvectors of the dependent variable may fail to fully control for phylogenetic structure when the predictor also contributes overlapping or unique phylogenetic signals. We therefore recommend using the union of eigenvectors from both traits (*EV*_*U*) in PVR models to more effectively remove shared phylogenetic signal and improve correlation accuracy.

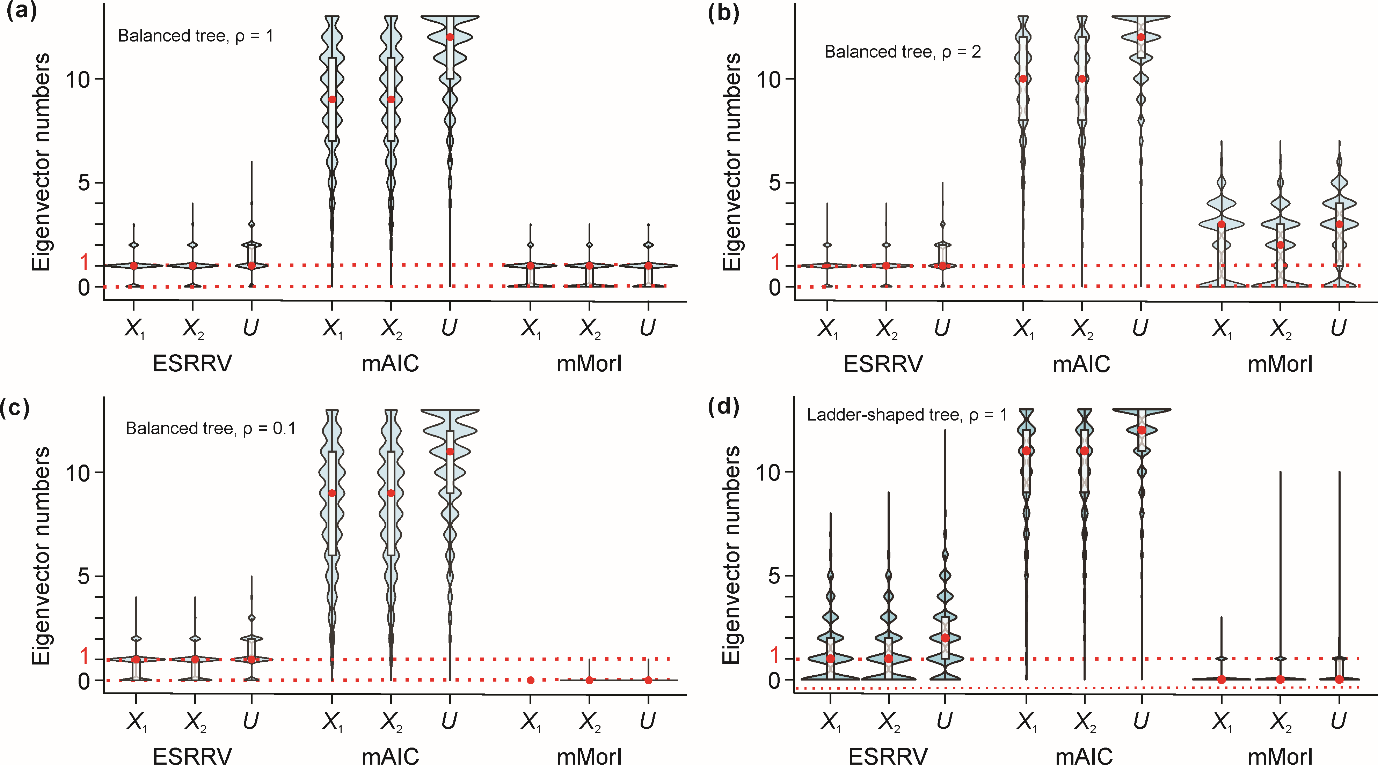

Fig. S1. Eigenvectors retained by three selection criteria (ESRRV, mAIC and mMorI) for a 16-species phylogenetic tree. EVs were picked separately for *X*_1_ and *X2*; *EV_U* is their union. Violin outlines depict the full distribution of selected eigenvector counts; white boxes show the inter-quartile range, black whiskers the 5th–95th percentiles, and red dots mark the median of each distribution. The red dashed line marks the ~1 predictor upper bound implied by the conventional samples‐per‐variable ≥ 10 guideline, calculated here assuming one non‐EV predictor. Selection procedures are described in Materials and Methods; the data used to plot this figure are accessible at https://github.com/zhenglinchen/PVR/.

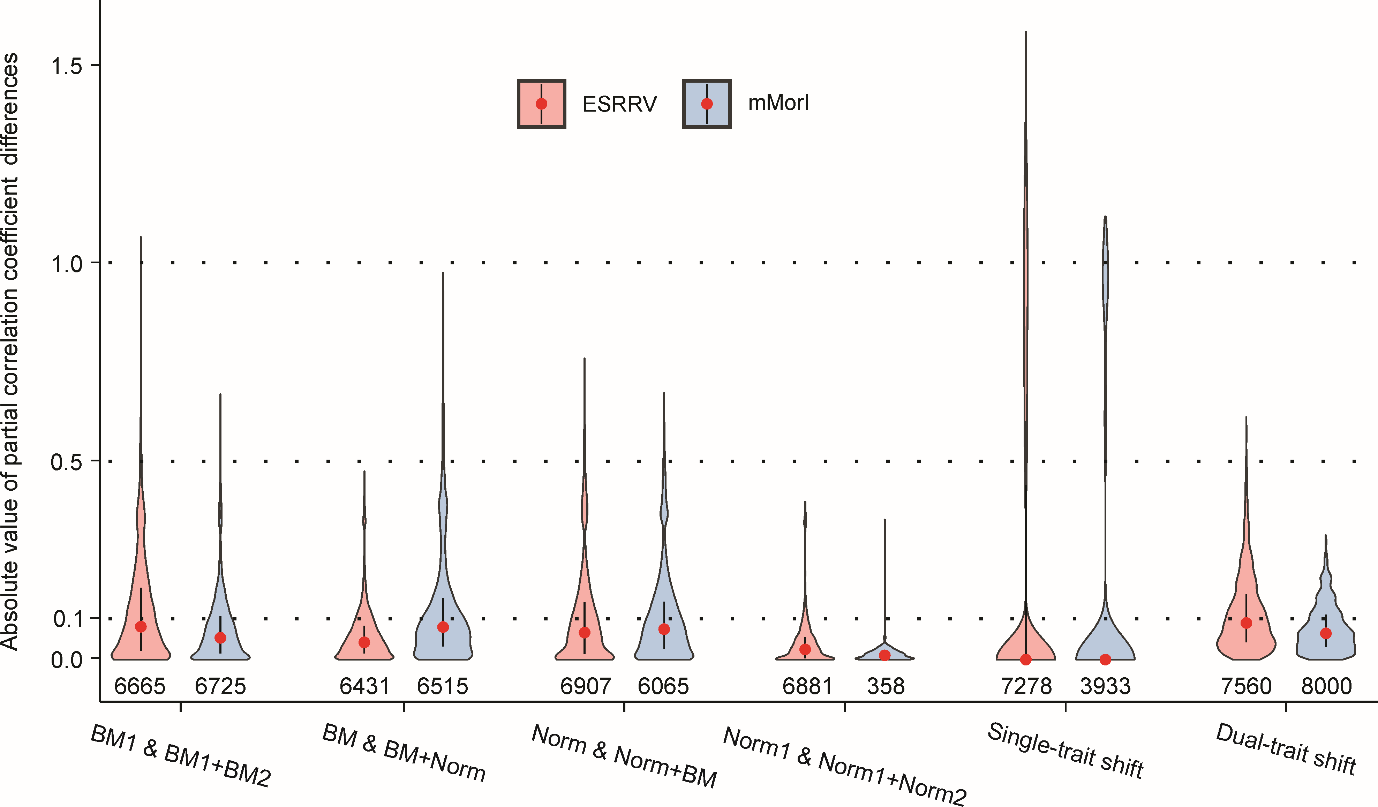

Fig. S2. Absolute differences in partial-correlation estimates ∣Δρ∣ between PVR models that use eigenvectors from *X*_1_ (*EV*_*X*_1_) or *X*_2_ (*EV*_*X*_2_) when the two models reach the *same* conclusion on significance (*p* < 0.05 vs ≥ 0.05) and slope sign. Violin colours correspond to the EV-selection methods (ESRRV, mMorI). Each violin shows the kernel-density distribution of |pcor(*X*_1_, *X*_2_ + *EV*_*X*_1_) − pcor(*X*_2_, *X*_1_ + *EV*_*X*_2_)|; numbers beneath indicate sample size (*n*). Red dots mark medians, black bars span the inter-quartile range, and thin whiskers extend to the 5th–95th percentiles. Dashed horizontal lines at ∣Δρ∣ = 0.1 and 0.5 denote "noticeable" and "large" effect-size thresholds, respectively (theoretical maximum = 2). Paired t-tests revealed significant differences in partial-correlation estimates between *EV*_*X*_1_ and *EV*_*X*_2_ in every scenario (all *p* < 0.001) except when mMorI was applied to the Norm1 & Norm1+Norm2 scenario data, where the test was non-significant (*n* = 358, *p* = 0.375). Full simulation and EV-selection details are given in *Materials and Methods*; the data used to plot this figure are accessible at https://github.com/zhenglinchen/PVR/.

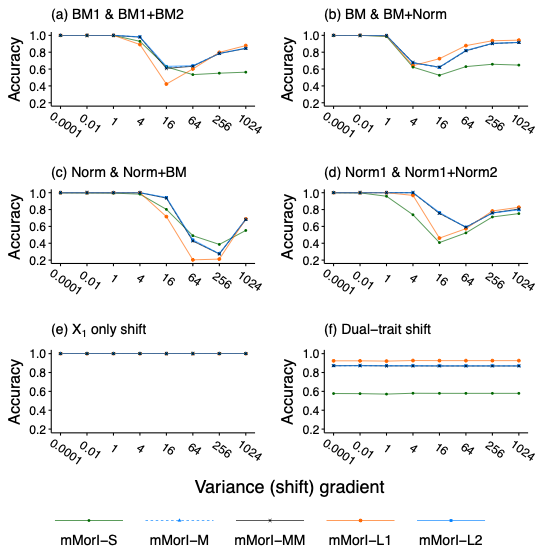

Fig. S3. Performance of five estimators within the EV_U-based PVR framework on 128-species balanced phylogenetic trees (ρ = 1). Comparisons were conducted under six evolutionary scenarios: (a) BM1 & BM1+BM2; (b) BM & BM+Norm; (c) Norm & Norm+BM; (d) Norm1 & Norm1+Norm2; (e) *X*_1_-only shift; and (f) Dual-trait shift. All eigenvectors (EVs) were selected using the mMorI criterion. For details on mMorI, scenario definitions, and accuracy measures, see Materials and Methods. The data used to plot this figure are accessible at https://github.com/zhenglinchen/PVR/.

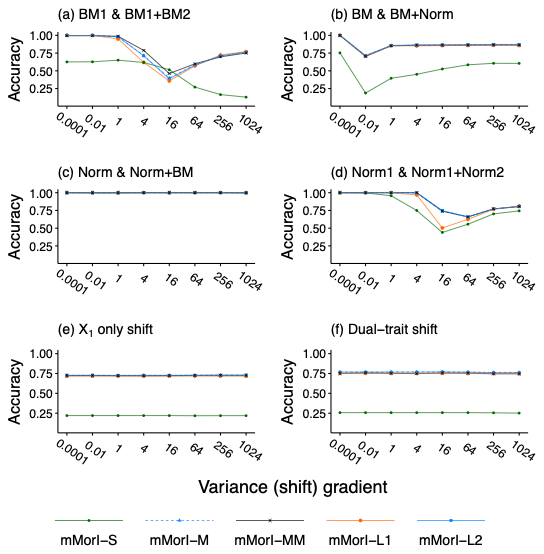

Fig. S4. Performance of five estimators within the EV_U-based PVR framework on 128-species balanced phylogenetic trees (ρ = 2). Comparisons were conducted under six evolutionary scenarios: (a) BM1 & BM1+BM2; (b) BM & BM+Norm; (c) Norm & Norm+BM; (d) Norm1 & Norm1+Norm2; (e) X1-only shift; and (f) Dual-trait shift. All eigenvectors (EVs) were selected using the mMorI criterion. For details on mMorI, scenario definitions, and accuracy measures, see Materials and Methods. The data used to plot this figure are accessible at https://github.com/zhenglinchen/PVR/.

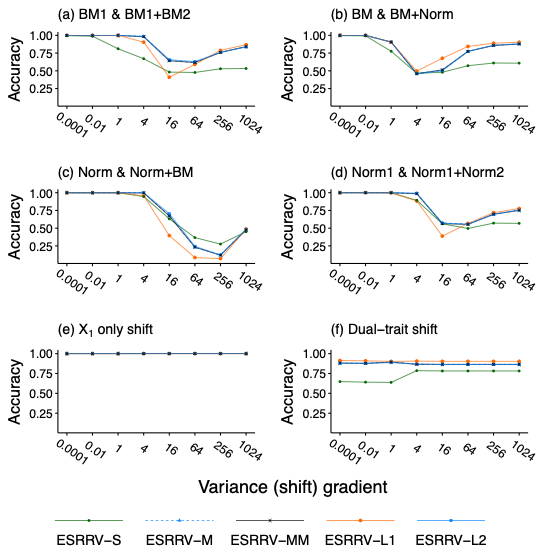

Fig. S5. Performance of five estimators within the EV_U-based PVR framework on 128-species balanced phylogenetic trees (ρ = 0.1). Comparisons were conducted under six evolutionary scenarios: (a) BM1 & BM1+BM2; (b) BM & BM+Norm; (c) Norm & Norm+BM; (d) Norm1 & Norm1+Norm2; (e) X1-only shift; and (f) Dual-trait shift. All eigenvectors (EVs) were selected using the ESRRV criterion. For details on ESRRV, scenario definitions, and accuracy measures, see Materials and Methods. The data used to plot this figure are accessible at https://github.com/zhenglinchen/PVR/.

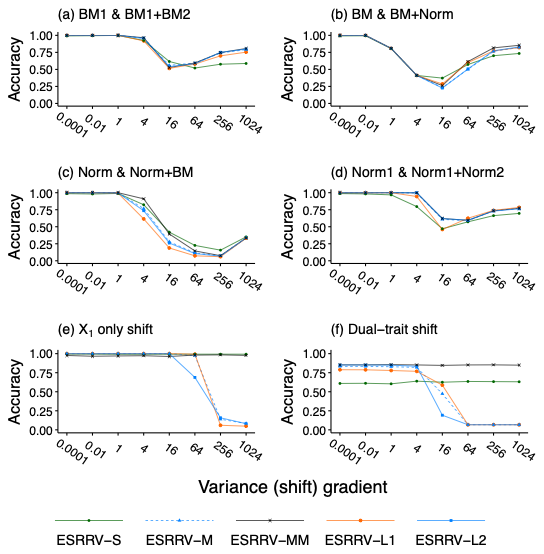

Fig. S6. Performance of five estimators within the EV_U-based PVR framework on 128-species ladder-shaped phylogenetic trees (ρ = 1). Comparisons were conducted under six evolutionary scenarios: (a) BM1 & BM1+BM2; (b) BM & BM+Norm; (c) Norm & Norm+BM; (d) Norm1 & Norm1+Norm2; (e) X1-only shift; and (f) Dual-trait shift. All eigenvectors (EVs) were selected using the ESRRV criterion. For details on ESRRV, scenario definitions, and accuracy measures, see Materials and Methods. The data used to plot this figure are accessible at https://github.com/zhenglinchen/PVR/.

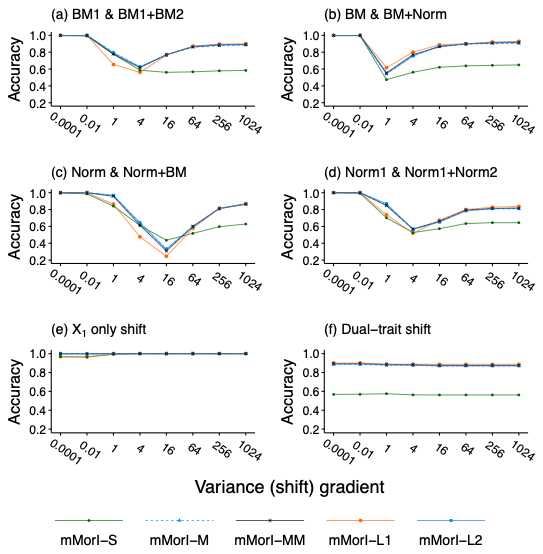

Fig. S7. Performance of five estimators within the EV_U-based PVR framework on 16-species balanced phylogenetic trees (ρ = 1). Comparisons were conducted under six evolutionary scenarios: (a) BM1 & BM1+BM2; (b) BM & BM+Norm; (c) Norm & Norm+BM; (d) Norm1 & Norm1+Norm2; (e) X1-only shift; and (f) Dual-trait shift. All eigenvectors (EVs) were selected using the mMorI criterion. For details on mMorI, scenario definitions, and accuracy measures, see Materials and Methods. The data used to plot this figure are accessible at https://github.com/zhenglinchen/PVR/.

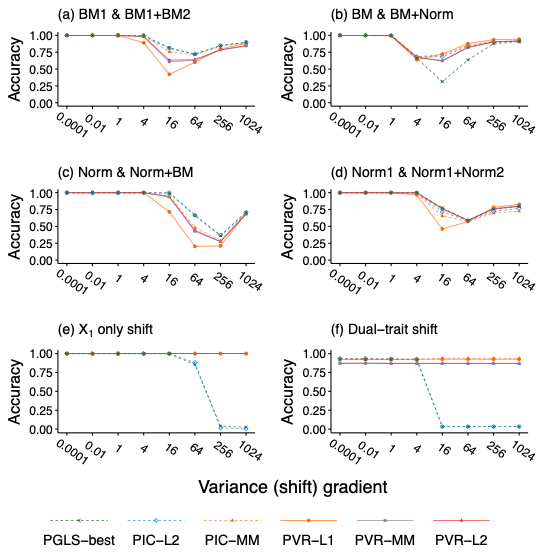

Fig. S8. Performance of PVR, PGLS, and PIC on 128-species balanced phylogenetic trees (ρ = 1). Comparisons were conducted under six evolutionary scenarios: (a) BM1 & BM1+BM2; (b) BM & BM+Norm; (c) Norm & Norm+BM; (d) Norm1 & Norm1+Norm2; (e) X1-only shift; and (f) Dual-trait shift. All eigenvectors (EVs) were selected using the mMorI criterion. For details on mMorI, scenario definitions, and accuracy measures, see Materials and Methods. The data used to plot this figure are accessible at https://github.com/zhenglinchen/PVR/.

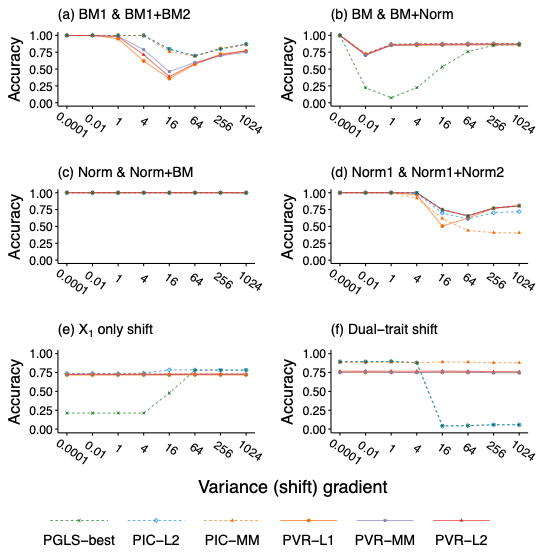

Fig. S9. Performance of PVR, PGLS, and PIC on 128-species balanced phylogenetic trees (ρ = 2). Comparisons were conducted under six evolutionary scenarios: (a) BM1 & BM1+BM2; (b) BM & BM+Norm; (c) Norm & Norm+BM; (d) Norm1 & Norm1+Norm2; (e) X1-only shift; and (f) Dual-trait shift. All eigenvectors (EVs) were selected using the mMorI criterion. For details on mMorI, scenario definitions, and accuracy measures, see Materials and Methods. The data used to plot this figure are accessible at https://github.com/zhenglinchen/PVR/.

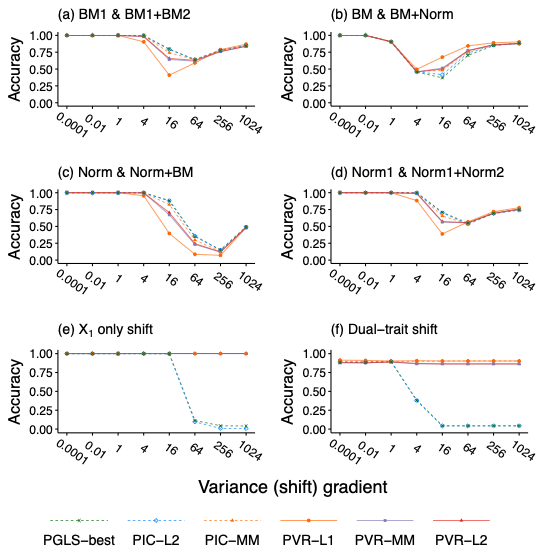

Fig. S10. Performance of PVR, PGLS, and PIC on 128-species balanced phylogenetic trees (ρ = 0.1). Comparisons were conducted under six evolutionary scenarios: (a) BM1 & BM1+BM2; (b) BM & BM+Norm; (c) Norm & Norm+BM; (d) Norm1 & Norm1+Norm2; (e) X1-only shift; and (f) Dual-trait shift. All eigenvectors (EVs) were selected using the ESRRV criterion. For details on ESRRV, scenario definitions, and accuracy measures, see Materials and Methods. The data used to plot this figure are accessible at https://github.com/zhenglinchen/PVR/.

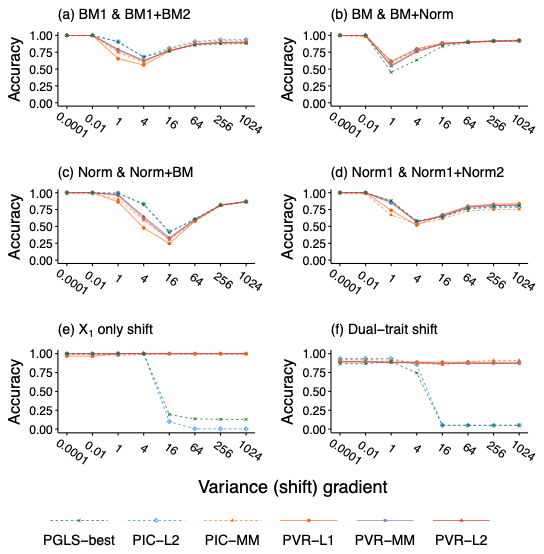

Fig. S11. Performance of PVR, PGLS, and PIC on 16-species balanced phylogenetic trees (ρ = 1). Comparisons were conducted under six evolutionary scenarios: (a) BM1 & BM1+BM2; (b) BM & BM+Norm; (c) Norm & Norm+BM; (d) Norm1 & Norm1+Norm2; (e) X1-only shift; and (f) Dual-trait shift. All eigenvectors (EVs) were selected using the mMorI criterion. For details on mMorI, scenario definitions, and accuracy measures, see Materials and Methods. The data used to plot this figure are accessible at https://github.com/zhenglinchen/PVR/.
